# A Unified Structure-based Deep Learning Framework for High-Throughput Screening of Protein-Binding RNAs

**DOI:** 10.64898/2026.09.07.749737

**Authors:** Yihao Zhao, Jing Han, Jike Wang, Jianfeng Chu, Yu Kang, Tingjun Hou

**Affiliations:** College of Pharmaceutical Sciences, Zhejiang University, Hangzhou 310058, Zhejiang, China; Institute of Materia Medica, Fujian Academy of Chinese Medical Science, Fuzhou 350003, Fujian, China; Zhejiang Provincial Key Laboratory for Intelligent Drug Discovery and Development, Jinhua 321016, Zhejiang, China

**Keywords:** protein-RNA interactions, machine learning, deep learning, virtual screening

## Abstract

Protein-RNA interactions regulate diverse biological processes and are increasingly exploited in therapeutic RNA discovery, but accurate inferences of nucleotide preferences and reliable structure prediction remain challenging. Here, we present PRIS, a unified structure-based deep-learning framework that combines two complementary components: PRISeq for nucleotide probability estimation at each RNA position and PRIScore for residue-nucleotide distance prediction to discriminate native-like from incorrect poses. Both share a feature extractor that integrates an Anti-Symmetric Graph Attention Network (A-GAT) with sparse k-Maximum Inner Product (k-MPI) attention to capture long-range interactions across large graphs. PRIScore improves the selection of native-like protein-RNA predictions generated by AlphaFold3, achieving a top-1 success rate of 81.91% on a docking benchmark, compared to 79.26% for AlphaFold3. The selected structures are then fed into PRISeq, which infers position-specific binding preferences and screens RNA libraries. On a PWM benchmark, PRISeq achieved a mean absolute error (MAE) of 0.75, outperforming FoldX, Rosetta-based scoring functions, and NA-MPNN. In virtual screening against MS2 protein, PRISeq screens 129,248 RNA hairpins within 11.95 seconds, achieving the highest EF_0.5%_ of 14.40, approximately double the best baseline. PRIS also effectively enriches active aptamers against NELF-E and GFP while preserving sequence diversity. By integrating structure selection with binding-preference inference, PRIS provides an efficient framework for large-scale RNA library screening and aptamer design.

## Introduction

Protein-RNA interactions (PRIs) regulate nearly every stage of RNA metabolism, including transcription, splicing, localization, translation, and degradation^1, 2^. PRIs are also related to viral replication, host defense, and pathogenesis of various diseases^3–5^. In addition to their fundamental biological roles, PRIs are increasingly exploited in therapeutic RNA discovery^6^. Synthetic RNAs engineered for high affinity to proteins are being developed for various applications, including diagnostics, molecular imaging, biomarker discovery, target validation, therapeutics, and drug delivery^7^. The rational design of such protein-binding RNAs (PBRs) requires accurate inferences of protein–RNA binding preferences based on accurate protein–RNA complex structures.

Regarding the inferences of protein–RNA binding preferences, a central requirement is to determine not only whether a protein binds RNA, but also which nucleotides it prefers at each position and how sequence variation alters binding. Such preferences are usually represented by position weight matrices (PWMs), which specify the probability of finding each of the four nucleotides at each position in the RNA-binding protein (RBP) binding site^8^. The PWM information can be experimentally obtained through RNA Bind-n-Seq (RBNS)^9^, sequencing and sequence specificity landscapes (SEQRS)^10^, RNAcompete-S^11^, systematic evolution of ligands by exponential enrichment combined with high-throughput sequencing (SELEX-seq)^12^, and high-throughput RNA-SELEX (HTR-SELEX)^13^. However, these experiments require extensive synthesis, selection, and quantitative binding measurements^14, 15^, limiting the number of proteins and sequence spaces that can be examined. Computational methods that infer RNA-binding preferences from structural information could therefore accelerate the annotation of RBPs and the discovery of functional RNA ligands.

Existing computational approaches provide complementary but incomplete solutions. Sequence-based methods completely ignore the structural information. The inherent flexibility and diverse conformations of nucleic acids complicate accurate inference of protein–RNA binding preferences using sequence-based methods. Consequently, structure-based methods are generally expected to yield higher inference accuracy compared to sequence-based methods^16^. Structure-based empirical scoring functions (SFs), including FoldX^17, 18^ and Rosetta energy functions^19, 20^, are sensitive to the input conformation. However, their computational cost becomes prohibitive as RNA length and library size increase in high-throughput virtual screening. A recent deep learning (DL)-based sequence design model, NA-MPNN^21^, offers accurate inferences of DNA-binding PWM, but its ability to infer RNA-binding PWMs was not evaluated in the original study^21^. Consequently, there remains a need for an efficient structure-based computational method capable of inferring RNA-binding preferences from a single protein-RNA complex structure.

It is also necessary to obtain accurate protein–RNA complex structures for the design of PBRs. The number of experimentally determined protein–RNA complex structures is very limited^22^. Due to the inherent conformational flexibility of proteins and RNAs, many weak and transient PRIs cannot be captured by experimental techniques, such as X-ray crystallography, nuclear magnetic resonance (NMR) and cryogenic electron microscopy (cryo-EM)^23^. Recent DL-based structure prediction methods, such as AlphaFold3^24^, have expanded access to protein-RNA structure predictions, but the conformational flexibility of RNA makes pose selection difficult. Consequently, even when the protein structure is accurately predicted, the RNA may be mispositioned relative to the protein or even with an incorrect secondary structure. Such errors can then propagate into sequence design or virtual screening. A common strategy is to generate many alternative structure predictions and then use a SF to select the best^25^. This strategy requires a reliable scoring function capable of distinguishing native-like binding poses from incorrect ones.

Methodological advances in both protein-RNA binding preference inference and protein-RNA complex structure prediction are important for the rational design of PBRs. Existing methods typically address these two tasks in isolation. Preference inference methods generally assume that the input structures are reliable and cannot assess their quality, whereas structure prediction methods do not provide the position-specific nucleotide preferences required for PBR sequence design. Manual examination is often required to correlate molecular interaction details from structural data with binding specificity data^26, 27^. Therefore, integrating the corresponding computational tools into a unified framework is essential.

Here, we developed PRIS, a unified structure-based framework comprising two DL-based components, PRISeq and PRIScore. Both components were introduced in this study and adopted the same feature extraction module but differed in their probability estimation modules. Their feature extractor combined an Anti-Symmetric Graph Attention Network (A-GAT), which incorporated graph attention^28^ into the stable and non-dissipative propagation framework of Anti-Symmetric Deep Graph Network (A-DGN)^29^, with k-Maximum Inner Product (k-MIP)^30^, which yielded a sparse and flexible attention pattern that approximated full attention. The former preserved long-range dependencies between nodes across multiple layers and the latter reduced computational cost when processing large graphs. Regarding probability estimation modules, PRISeq estimated probability distributions of nucleotide types, whereas PRIScore estimated probability distributions of residue-nucleotide distances. In the PRIS workflow, PRIScore was used to improve the selection of native-like protein-RNA predictions if the crystal structure was unavailable. The selected prediction was then input to PRISeq to infer RNA-binding preferences or screen candidate RNA sequences.

We curated an extensive dataset of 9,110 protein-nucleic acid complex structures sourced from Protein Data Bank (PDB)^31^ for model training. To comprehensively evaluate existing PRIS, we also compiled three validation sets, including 1) a PWM dataset that comprises 15 crystal structures of protein-RNA complexes with experimentally determined RNA-binding PWMs, involving 108 nucleotide positions^14, 32^, 2) a docking dataset that comprises 188 crystal structures, which are directly retrieved from three published protein-RNA docking benchmarks^33–35^, and 3) a screening dataset that comprises 3 target proteins comprised three target proteins, including MS2, NELF-E, and GFP. For each specific target protein, the binding affinities of equal-length RNA sequences are measured by the same research group using the same bioassay. The MS2 ligand library comprises 129,248 RNA hairpin sequences^36^. The NELF-E and GFP ligand libraries comprise 9,833 and 1,875 RNA aptamer sequences, respectively^37^. PRIS is rigorously tested on the PWM, docking and screening datasets, demonstrating its ability to infer RNA-binding preferences, identify native-like binding predictions, and prioritize high-affinity RNA ligands. By connecting predictions selection with preference inference, PRIS provides an efficient route from experimental or predicted protein-RNA structures to large-scale RNA screening and RNA aptamer design.

## Materials and Methods

### Dataset preparation

A total of four datasets, i.e., the structure, PWM, docking, and screening datasets, were curated in this study. The structure dataset was used for model training. The remaining three sets were used as the test sets for model evaluation. The structure dataset consisted of 9,110 protein-nucleic acid complex structures retrieved from PDB (before April 5, 2026). Among them, 328 protein-RNA complexes were labeled with affinity data from the PDBbind database (v2020)^38^ or the PRA310 dataset^39^ to fine-tune the model, forming the binding affinity dataset PRA328.

The PWM dataset comprised 15 crystal structures of protein-RNA complexes with experimentally determined RNA-binding PWMs. The crystal structures were collected from PDB and the PWMs were collected from CISBP-RNA dataset^14^ and SpliceAid-F dataset^32^. An ungapped alignment was performed between the sequences extracted from PDB and the sequences obtained from the CISBP-RNA and SpliceAid-F datasets.

The docking dataset comprised 188 crystal structures, which were directly retrieved from three published protein-RNA docking benchmarks^33–35^. The duplicate crystal structures were identified based on their PDB IDs and removed, with only one instance of each structure retained. In addition, only crystal structures containing standard amino acid residues and nucleotides were retained, and their sequences were used as inputs to AlphaFold3.

The screening dataset comprised three target proteins, including MS2, NELF-E, and GFP. For each specific target protein, the binding affinities of equal-length RNA sequences were measured by the same research group using the same bioassay. The MS2 ligand library comprised 129,248 RNA hairpin sequences^36^. The NELF-E and GFP ligand libraries comprised 9,833 and 1,875 RNA aptamer sequences, respectively^37^. The top 1% RNAs ranked by experimental binding data were regarded as the active ligands for a specific protein. The others were regarded as the decoys.

### PRIS

The model architecture of PRIS is depicted in **Figure 1**, which consists of graph representation, feature extraction, and probability estimation modules. PRISeq and PRIScore are two branches of PRIS. They adopt the same protein graph representations and feature extraction modules but differ in their probability estimation modules. PRISeq estimates probability distributions of nucleotide types, whereas PRIScore estimates probability distributions of residue-nucleotide distances. The architecture of PRISeq is described in the following sections.

**Figure 1.**
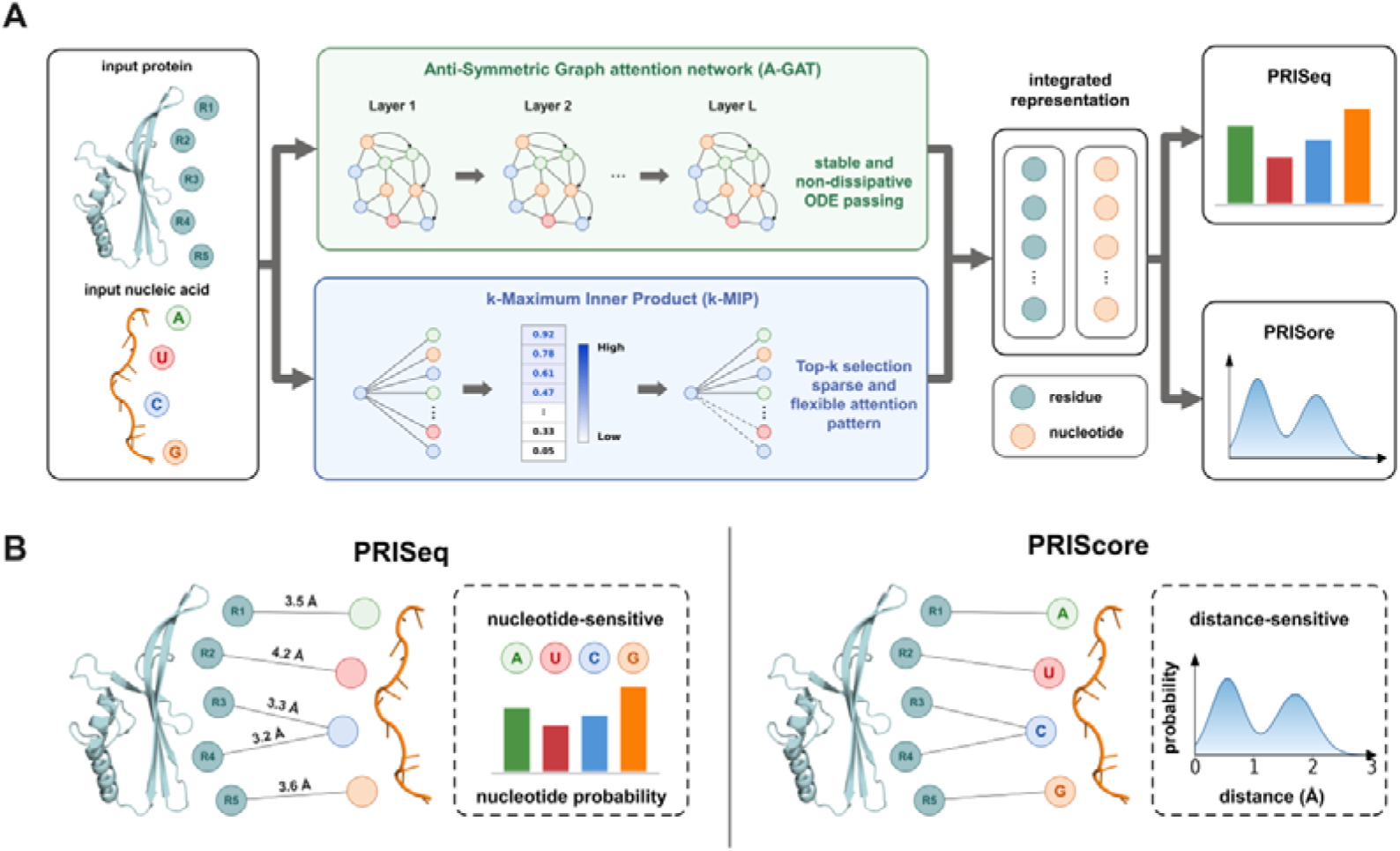
Overview of the PRIS framework. (A) Schematic illustration of the PRIS framework. Given a protein-RNA complex structure, residue and nucleotide features are extracted and processed through two complementary modules, A-GAT and k-MIP. A-GAT preserves long-range dependencies between nodes across multiple layers through stable and non-dissipative graph ODEs. k-MIP selects the most relevant key nodes for each query via a top-k operation, yielding a sparse and flexible attention that approximates full attention. Their learned representations are integrated and passed to the two task-specific branches of PRIS, PRISeq and PRIScore. (B) PRISeq estimates probability distributions of nucleotide types, whereas PRIScore estimates probability distributions of residue-nucleotide distances.

### Graph representations

In the graph representation module, the graph for each nucleic acid was constructed at the nucleotide level, which was demonstrated to be effective in our previous study^40^. In this graph, nodes represented the nucleotides of the entire nucleic acid, and edges represented the interactions between any two nucleotides with a maximum distance of 10.0 Å. To enable the prediction of general RNA-binding preferences rather than memorization of sequence-specific patterns, the base identities and base-specific atomic details were removed. Only the sugar-phosphate backbone atoms (O3’, C3’, C4’, C5’, O5’, P, C2’, C1’, and O4’) were retained for each nucleotide. **Table S1** summarizes the features of each nucleic acid graph. Specifically, the node features include the self-distances and dihedral angles of each nucleotide, whereas the edge features include bonding states, C5’-to-C5’ distances, center-to-center distances, and the maximum and minimum distances between any two nucleotides.Similarly, the graph for each pocket was constructed at the residue level. The the binding pocket. Each pocket was then represented as a graph (*G^P^* = (*N^P^*, *E^P^*)), residues located within 10.0 Å radius around the co-crystallized RNA were defined as with nodes representing the residues in a pocket and edges representing the interactions between any two residues with a minimum distance of less than 10.0 Å. To incorporate sequence-derived contextual information, the corresponding protein sequence was processed using ESM3^41^, and the residue-level hidden representations generated by the pretrained model were aligned with the respective graph nodes and incorporated into their features. The features of protein graph are outlined in **Table S2**. The node features include amino acid type, self-distances, dihedral angles and the hidden representations from ESM3 for each residue, and the edge features include bonding states, CA-to-CA distances, center-to-center distances, and the maximum and minimum distances between any two residues.

### Feature extraction

The protein and nucleic acid graphs shared the same model architecture but employed independent feature extractors to transform the features into their respective hidden representations. For a graph with node features 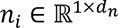 for node i and edge features 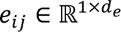 for the edge between node i and its neighboring node j, these features were first embedded into d-dimensional initial hidden representations by linear transformations:

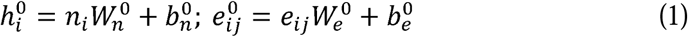

where 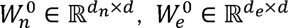, and 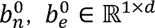, are the weights and biases of the linear layers, respectively; 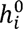 and 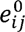 are the node and edge features in the first hidden layer, respectively.

In each layer of the feature extraction module, the hidden representation was processed through an A-DGN, a framework for effective long-term information propagation in DGN architectures. By applying a stable and non-dissipative ordinary differential equation (ODE) system on graphs, A-DGN preserved long-term dependencies between nodes and prevented gradient explosion and vanishing. GATs assigned learnable attention coefficients to neighboring nodes, enabling the adaptive aggregation of information from structurally and functionally relevant interactions. To combine this adaptive neighborhood weighting with the stable and non-dissipative properties of A-DGN, we incorporated GAT as the message-passing function within the A-DGN framework and termed the resulting architecture A-GAT. The A-GAT operation of the *l*th layer was described as follows:

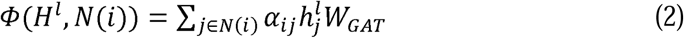

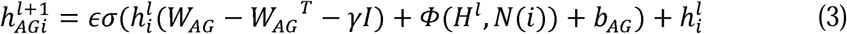

where 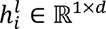 is the hidden representation of node i, 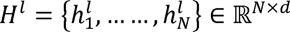 is the set of 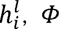 is the GAT function that aggregates nodes information, *N*(*i*) denotes the out-neighbors set of node i, *α_ij_* is the normalized attention coefficient, and *W_GAT_* ɛ ℝ*^d×d^* is the corresponding input linear transformation’s weight matrix in GAT; ɛ is the discretization step, 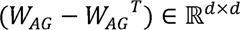 is the anti-symmetric weight matrix, γ is a hyper-parameter that regulates the strength of the diffusion, *I* ɛ ℝ*^d×d^* is the identity matrix, and *b_AG_* ɛ ℝ*^d×d^* is the bias vector in A-GAT.

A-GAT enabled stable and non-dissipative message passing across multiple layers, making it suitable for deep architectures. However, its reliance on local message passing limited its ability to capture global interactions, particularly in large graphs, where long-range dependencies became increasingly important. To address this limitation, we further incorporated the k-MIP attention, which selected the most relevant key nodes for each query through a top-k operation, yielding a sparse yet flexible attention pattern. Combined with symbolic-matrix-based attention score computation, k-MIP achieved linear memory complexity and practical speedups of up to an order of magnitude over all-to-all attention without compromising the expressive power of graph transformers. The k-MIP operation of the *l*th layer was described as follows:

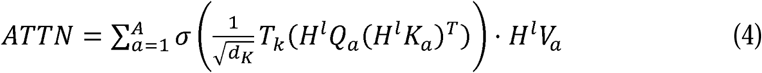

where *T_K_* is the top-k operation that retains only the k largest elements in each row and sets all other elements to -∞; *Q_α_*, *K_α_* ɛ ℝ*^d×m^* and *V_α_* ɛ ℝ*^d×d^* are the *α*th weights from *A*-sets of Query, Key and Value weight matrices, respectively; *σ* represents the softmax operation.

The outputs of A-GAT and k-MIP were independently subjected to dropout, residual connections, and batch normalization. The resulting representations were then aggregated and passed through a 2-layer multilayer perceptron (MLP). This hybrid architecture combined the capacity of A-GAT for stable propagation through deep networks with the scalability of k-MIP for integrating information across large graphs. The MLP operation of the th layer was described as follows:

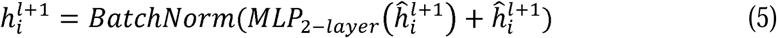

where 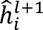 and 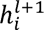 denote the aggregated and final node representations in the (*l* + 1) th layer of feature extraction module, respectively; *BatchNorm* denotes the batch normalization operation. The output 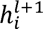 was passed through the subsequent layers, and the output of the final layer was fed into the nucleotide probability estimation module.

### Nucleotide probability estimation

In the nucleotide probability estimation module, the extracted residue and nucleotide node representations were integrated to estimate the probability distributions of nucleotide types. Residue-nucleotide distance information was additionally incorporated as an auxiliary geometric feature. The distance features *D_x,y_* were first processed using an MLP and then combined with the integrated node representations. The resulting pairwise features were passed through a second MLP, followed by a linear transformation and an exponential linear unit (ELU) activation, to quantify the contribution of residue x to the nucleotide probability distribution *Z_x,y_* ɛ ℝ^1×12^ of nucleotide y. The contributions from all residues interacting with nucleotide y were summed. In addition, the intrinsic contribution of nucleotide y was estimated from its node representation 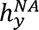 and added to the sum of the residue contributions. The corresponding operations were described as follows:

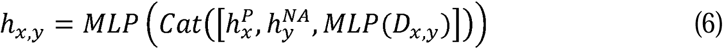

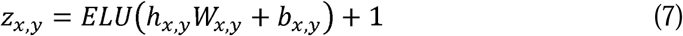

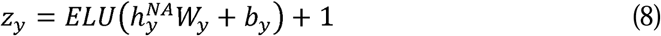

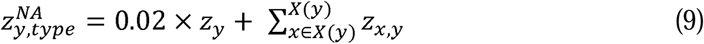

where 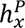 and 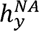 denote the extracted residue and nucleotide node representations, respectively; *D_x,y_* denotes residue-nucleotide distance features, calculated using all backbone and side-chain atoms of the residue x and only the backbone atoms of the nucleotide y; *Cat* represents the concatenation operation; *ELU* is a type of activation function; *W_x,y_* ɛ ℝ*^4d×d^*, *W_y_* ɛ ℝ*^d×d^*, and *b_x,y_*, *b_y_* ɛ ℝ^1*×d*^ are the weights and biases of the linear transformation, respectively; *X*(*y*) denotes the set of residues interacting with nucleotide y; 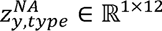 denotes the predicted 12-dimensional logit vector, with each dimension corresponding to one of the 12 nucleotide types: A, G, C, U, I, N, DA, DG, DC, DT, DU and DI. Applying the softmax function to 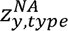 yields the probability distribution of nucleotide types.

### Loss function

The total loss *L* was composed of cross-entropy losses *L_nt_* and *L_bt_* and correlation loss *L_affi_*. The cross-entropy losses *L_nt_* and *L_bt_* supervised the predictions of nucleotide types and their bond types, respectively. The Pearson correlation coefficient (PCC) between the predicted and experimental binding affinities of a batch of protein-nucleic acid complexes was calculated and shared as the correlation loss *L_affi_*. The loss function was defined as follows:

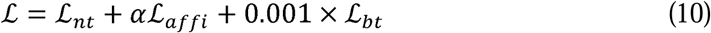

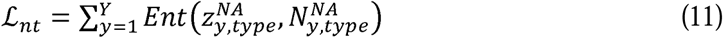

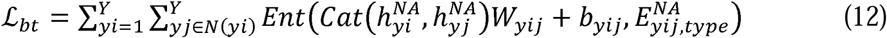

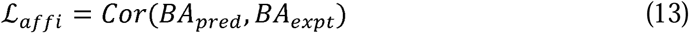

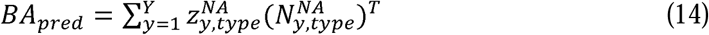

where α is the weight of 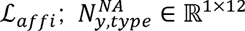 denotes the one-hot encoding of the nucleotide type at nucleotide y; *Ent* stands for the cross-entropy operation; 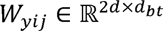 and 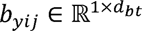 are the weight and bias of the linear transformation, respectively; 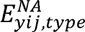 represents the type of bond between nucleotide yi and nucleotide yj; *Cor* represents the PCC operation; *BA_pred_* and *BA_expt_* represent the predicted and experimental binding affinities, respectively.

### Comparison of PRISeq and PRIScore

As mentioned, PRISeq and PRIScore used the same protein graph representations and feature extraction modules but differed in their nucleic acid graph representations and probability estimation modules. The nucleic acid graph representation of PRIScore was the same as that introduced in our previous study, PDIScore^40^. By contrast, base identities and base-specific atomic details were removed from the nucleic acid graph representation of PRISeq to facilitate the prediction of general RNA-binding preferences rather than the memorization of sequence-specific patterns.

Regarding the probability estimation modules, PRIScore used a mixture density network (MDN) to estimate the residue-nucleotide distance distributions from the concatenated features, following the approach introduced in PDIScore^40^. The residue-nucleotide distance was treated as a continuous variable in PRIScore. By contrast, PRISeq estimated the probability distributions of nucleotide types rather than residue-nucleotide distances. As a discrete variable, nucleotide type could not be directly modeled by an MDN. Therefore, PRISeq employed a softmax layer to generate the nucleotide probability distributions.

### Model training

To prevent data leakage between the structure dataset and the test sets, their protein chains were jointly clustered based on sequence similarity using the program MMseqs2^42^. In the structure dataset, 32, 836, and 22 structures contained at least one protein chain sharing more than 40% sequence identity with a protein chain in the PWM, docking, and screening datasets, respectively. These structures were excluded from training.

Following the parameterization strategy in our previous study^40^, we trained the model with two steps: normal training and fine-tuning. During the initial training phase, *α* was set to 0, and the prepared structure dataset was randomly divided into a validation set of 700 complexes and a training set comprising the remaining complexes. In the subsequent fine-tuning phase, was adjusted to 0.5, and the PRA328 dataset was also randomly split into a validation set of 40 complexes and a training set comprising the remaining complexes.

### Model evaluation

PRIS was evaluated on the PWM, docking and screening datasets. In the PWM dataset, model performance was assessed using accuracy (ACC), Spearman correlation coefficient (SCC), mean squared error (MSE), and mean absolute error (MAE). ACC indicated the sequence recovery, which measured the proportion of RNA positions for which the most preferred nucleotide was correctly identified, whereas SCC quantified the agreement between the predicted and experimentally determined rankings of the four nucleotide types. MSE and MAE measured the differences between the predicted and experimental nucleotide probabilities, with MSE assigning greater weight to large deviations and MAE reflecting the average absolute deviation. As for the docking set, model performance was assessed using the success rate (SR), defined as a prediction being successful if at least one of the top-ranked poses had a root-mean-square deviation (RMSD) value of less than 6.0 Å from the native pose. Both RNA RMSD(R) and protein RMSD(P) were required to satisfy the condition. The atom P from RNA and atom CA from protein were used to calculate RMSD(R) and RMSD(P), respectively. Regarding the screening set, model performance was assessed using enrichment factor (EF) and area under the receiver operating characteristic curve (AUROC). EF was calculated as the average percentage of true RNA binders among the top-scoring RNA candidates (1%, 5%, or 10%). AUROC quantified the model’s ability to distinguish binders from non-binders.

### Baselines

In addition to PRIS, several other methods were included as the baselines for comparison. In the PWM dataset, we used a random baseline, three traditional empirical SFs, including FoldX, Rosetta with the ref2015 energy function (Rosetta#ref2015), and Rosetta with the RNP-ddG energy function (Rosetta#RNP-ddG^20^), and an RNA sequence design method, NA-MPNN^21^. In the docking dataset, we used AlphaFold3 as a pose prediction baseline and three SFs, including FoldX, Rosetta#ref2015, and Rosetta#RNP-ddG to re-score the prediction generated by Alphafold3. In the screening dataset, we used three SFs, including FoldX, Rosetta#ref2015, and Rosetta#RNP-ddG for virtual screening.

In the PWM dataset, the scores of these SFs were converted into PWMs using the Boltzmann formula 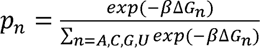 Starting from a crystal structure of the protein bound to its consensus RNA sequence, a structural ensemble was created by threading novel RNA sequences onto the binding site. Protein-RNA binding energies, Δ*G*, were then evaluated for each member of the structural ensemble. It was only necessary to compute Δ*G* values for all single-point mutations from the consensus-binding site^43^. We used Chimera^44^ to introduce single-point mutations and SFs to evaluate binding energies. Regarding NA-MPNN, we performed random autoregressive decoding 30 times at a temperature of 0.6, saved the per-position categorical probabilities from each run, and averaged them to obtain a predicted PWM for every RNA position, as described in the original study^21^. In the docking dataset, we used AlphaFold3 with random seeds ranging from 1 to 20 to generate 100 predictions for each complex. In the screening dataset, the virtual screening workflow using SFs involved several steps, including mutation generation, energy minimization, and energy evaluation. The reference crystal structures were first mutated using Chimera^44^ according to the target RNA sequences, followed by the minimization of the whole complexes using Rosetta^19^. The energy-minimized complexes were evaluated using the SFs to calculate their binding energies.

## Results and discussion

### PRISeq accurately infers RNA-binding preferences

In the PWM inference task, we evaluated the ability of PRISeq to infer the RNA-binding preferences of target proteins. Given a protein-RNA complex structure, PRISeq extracted and integrated residue and nucleotide features to predict the nucleotide probability distribution at each RNA position. The PWM dataset comprised 15 crystal structures of protein-RNA complexes with experimentally determined RNA-binding PWMs, involving 108 nucleotide positions. We compared PRISeq with a random baseline, three traditional empirical SFs, including FoldX, Rosetta#ref2015, and Rosetta#RNP-ddG, and an RNA sequence design method, NA-MPNN. The scores of these SFs were converted into PWMs using the Boltzmann formula, as detailed in Materials and Methods. Model performance was assessed using ACC, SCC, MSE, and MAE. ACC indicated the sequence recovery, which measured the proportion of RNA positions for which the most preferred nucleotide was correctly identified, whereas SCC quantified the agreement between the predicted and experimentally determined rankings of the four nucleotide types. MSE and MAE measured the differences between the predicted and experimental nucleotide probabilities, with MSE assigning greater weight to large deviations and MAE reflecting the average absolute deviation. As shown in **Table 1**, PRISeq consistently outperformed all baseline methods across the four evaluation metrics, achieving an ACC of 65.6%, an SCC of 0.43, an MSE of 0.43, and an MAE of 0.75. Compared with the best-performing baseline for each metric, PRISeq increased the ACC and SCC by 48.1% and 38.7%, respectively, and reduced the MSE and MAE by 23.2% and 25.7%, respectively. These results demonstrated that PRISeq effectively captured sequence-dependent differences in RNA-binding preferences within a shared structural context, providing a more accurate quantitative description of binding specificity than traditional empirical SFs.

**Table 1.** Performance of the methods in the PWM inference task.

| Methods | ACC(%) ↑ | SCC ↑ | MSE ↓ | MAE ↓ |
| --- | --- | --- | --- | --- |
| Random | 25.8 | 0.00 | 0.67 | 1.27 |
| FoldX | 32.7 | 0.02 | 0.66 | 1.17 |
| Rosetta#ref2015 | 42.1 | 0.13 | 0.58 | 1.02 |
| Rosetta#RNP-ddG | 44.3 | 0.21 | 0.65 | 1.01 |
| NA-MPNN | 42.9 | 0.31 | 0.56 | 1.04 |
| PRISeq | **65.6** | **0.43** | **0.43** | **0.75** |

We further examined how the performance of PRISeq varied with information content (IC), which was calculated using the equation^45^ *IC* = *2 +* Σ*_n_*_= *A,C,G,U,*_*P_n_log_2_P_n_*, where *P_n_* denotes the nucleotide probability derived from the PWM. The 108 nucleotide positions were classified into 77 highly conserved positions (experimental IC ≥ 1.0 bit) and 31 less conserved positions (experimental IC < 1.0 bit). The nucleotide distributions were more concentrated at highly conserved positions, reflecting strong preferences for specific nucleotides^46^. By contrast, the nucleotide distributions were broader nucleotide distributions at less conserved positions, showing greater degeneracy^47, 48^ (**Figure 2B**). PRISeq exhibited better performance and higher predicted IC at highly conserved positions (MAE = 0.74, predicted IC = 1.42) than at less conserved positions (MAE = 0.83, predicted IC = 1.20). Although PRISeq’s performance decreased at less conserved positions, its lower predicted IC reflected the degeneracy of these positions, thereby supporting the biological interpretability of the predicted PWMs.

**Figure 2.**
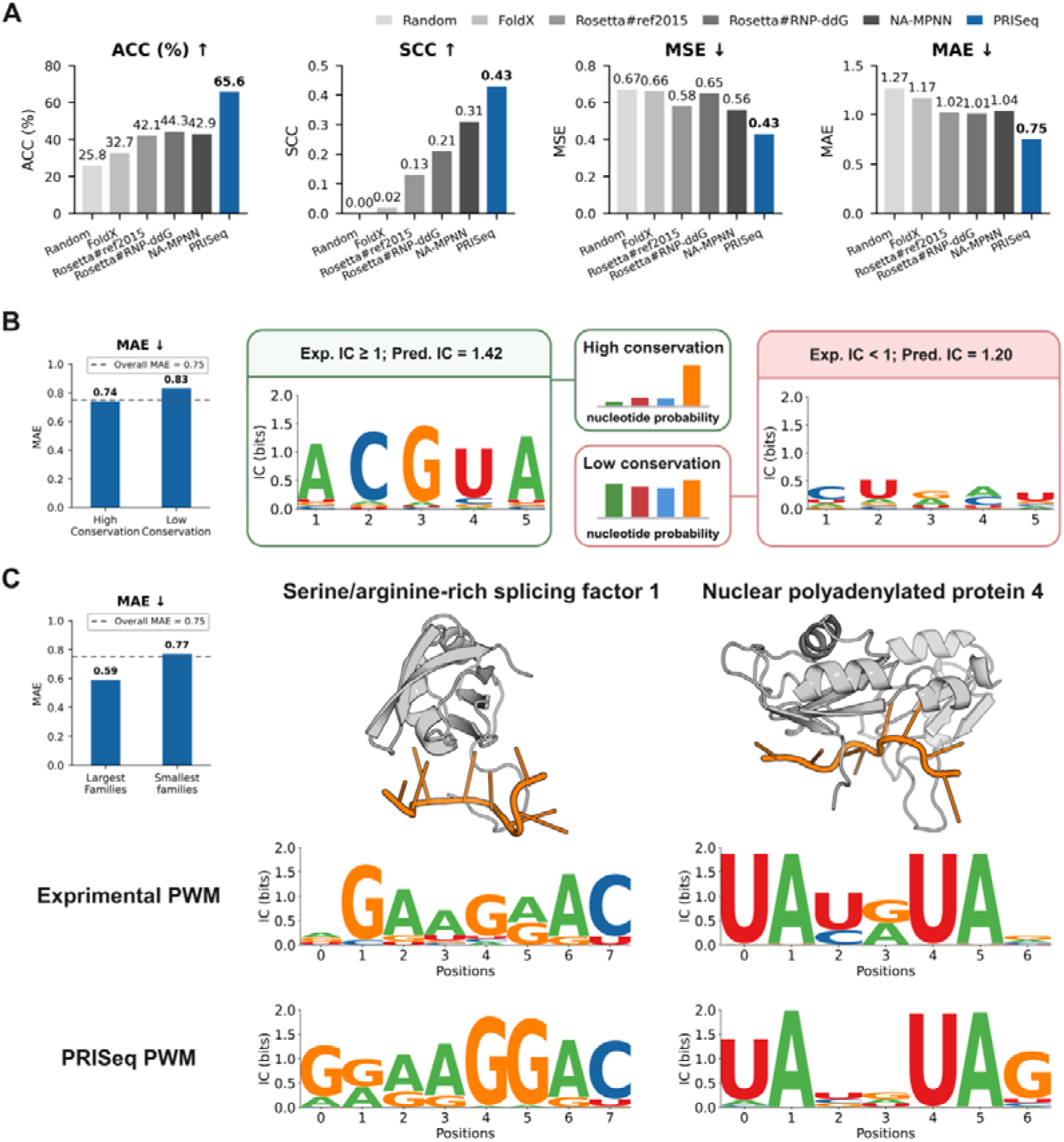
Performance of PRISeq in the PWM inference task. (A) Comparison of PRISeq with a random baseline, FoldX, Rosetta#ref2015, Rosetta#RNP-ddG, and NA-MPNN. Performance is evaluated in terms of ACC, SCC, MSE, and MAE. The upward and downward arrows indicate that larger and smaller values correspond to the better performance, respectively. (B) PRISeq performance at highly conserved (experimental IC ≥ 1 bit) and less conserved (experimental IC < 1 bit) nucleotide positions. The dashed line indicates the overall MAE of 0.75. Representative sequence logos illustrate more concentrated and broader nucleotide probability distributions at highly and less conserved positions, respectively. (C) PRISeq performance for protein families with the largest and smallest numbers of structures. Representative protein-RNA complex structures and comparisons between experimentally determined and PRISeq-predicted PWMs are shown for serine/arginine-rich splicing factor 1 and nuclear polyadenylated RNA-binding protein 4. Proteins and RNAs are shown in grey and orange, respectively. In the sequence logos, letter heights represent nucleotide IC in bits.

We also assessed the performance of PRISeq across protein families (**Figure 2C**). PRISeq achieved an average MAE of 0.59 for the families with the most structures. As an example, the MAE for serine/arginine-rich splicing factor 1 (SRSF1) was 0.51. SRSF1 is a key regulator of constitutive and alternative splicing^49^. According to the experimentally determined PWM, SRSF1 recognizes purine-rich sequence motifs and its binding promotes recognition of both constitutive and alternative exons during spliceosomal assembly^50^. PRISeq reproduced this purine-rich preference at most positions and 3’-terminal C preference, suggesting that its predicted PWM could recover sequence features associated with SRSF1-dependent exon recognition. The predicted PWM showed consistently low probabilities for nucleotide U at each position, suggesting that the U substitutions could weaken SRSF1 binding and consequently reduce splicing enhancer activity. This hypothesis was supported by previous mutational, in vitro splicing, and UV cross-linking/immunoprecipitation experiments^51^, showing the biological relevance of the predicted PWMs. Regarding the families with the fewest structures, PRISeq achieved an average MAE of 0.77. As an example, the MAE for Nuclear polyadenylated RNA-binding protein 4 (Nab4) was 0.30. Nab4 is a component of the cleavage and polyadenylation machinery that recognizes the UA-rich efficiency element and contributes to 3’ pre-mRNA processing and polyA cleavage site selection^52^. The predicted PWM was similar to the experimental PWM, showing the UA-rich motif recognized by Nab4. Notably, the average MAE for the families with the fewest structures (0.77) was close to the overall average MAE (0.75) with a relative deviation of 2.7%, suggesting that PRISeq learned intrinsic protein-RNA binding preferences rather than overfitting on family-specific patterns.

### PRISeq rapidly screens a 129,248-member RNA library against the MS2 protein

Besides PWM inference, we also applied PRISeq to rapidly screen a 129,248-member RNA library against the MS2 protein, which was involved in viral genome encapsidation through the interaction between a capsid protein dimer and the multiple packaging signals present in the RNA genome^53^. The binding affinities of 129,248 equal-length RNA hairpin sequences to the MS2 protein were experimentally measured^36^. The experimentally determined crystal structure (PDB ID: 1ZDK) of the MS2 protein bound to the wild-type RNA hairpin^54^ was used as input for PRISeq. We compared PRISeq with three SFs, including FoldX, Rosetta#ref2015, and Rosetta#RNP-ddG. Regarding these SFs, the virtual screening workflow involved several steps, including mutation generation, energy minimization, and energy evaluation, as detailed in Materials and Methods. Model performance was assessed using EF and AUROC. EF was calculated as the average percentage of true RNA binders among the top-scoring RNA candidates (0.5%, 1%, or 5%). AUROC quantified the model’s ability to distinguish binders from non-binders. As shown in **Table 2**, PRISeq consistently outperformed all baseline methods across the four evaluation metrics, achieving an EF_0.5%_ of 14.40, an EF_1%_ of 10.61, an EF_5%_ of 4.94, and an AUROC of 0.63. Compared with the best-performing baseline for EF, PRISeq approximately doubled EF_0.5%_ and EF_1%_, and approximately tripled EF_5%_. The improved performance across evaluation metrics demonstrated the potential of PRISeq for large-scale RNA screening.

**Table 2.** Performance of the methods in the virtual screening of a 129,248-member RNA library against the MS2 protein. Time denotes the average computation time per sequence using an NVIDIA A100 GPU.

| Methods | EF<sub>0.5%</sub> ↑ | EF<sub>1%</sub> ↑ | EF<sub>5%</sub> ↑ | AUROC ↑ | Time (s) ↓ |
| --- | --- | --- | --- | --- | --- |
| FoldX | 6.97 | 4.80 | 1.58 | 0.48 | 0.07 |
| Rosetta#ref2015 | 0.77 | 0.93 | 0.70 | 0.53 | 6.01 |
| Rosetta#RNP-dd<br>G | 0.46 | 0.62 | 1.52 | 0.61 | 5.42 |
| PRISeq | **14.40** | **10.61** | **4.94** | **0.63** | **9.25×10<sup>-5</sup>** |

PRISeq also achieved higher computational efficiency compared with the SFs. As described above, PRISeq required only one input structure, whereas the SFs had to evaluate large numbers of protein-RNA complex structures corresponding to different RNA hairpin sequences. PRISeq completed the virtual screening of a 129,248-member RNA library in a single run within 11.95 s. By contrast, FoldX, Rosetta#ref2015, and Rosetta#RNP-ddG required approximately 2.5 h, 9.0 days, and 8.1 days, respectively, to evaluate the same library. Moreover, the above runtimes excluded the time required for mutation generation and energy minimization, both of which would incur additional computational costs in the SF-based virtual screening workflow. On average, it took only 9.25×10^-5^ s for PRISeq to predict the binding affinity of each RNA hairpin sequence, achieving approximately three orders of magnitude acceleration compared with the fastest SF, FoldX (0.07 s). It should also be noted that the computational efficiency advantage of PRISeq became more pronounced as library size increased. PRISeq could screen the entire sequence space (4^19^ sequences) of the 19-nt RNA hairpin in a single run, achieving an average computation time of only 4.35×10^-11^ s. These results highlighted the capability of PRISeq to rapidly screen large RNA libraries.

In addition to achieving higher computational efficiency, PRISeq enriched for experimentally active sequences and preserved substantial sequence diversity (**Figure 3C**). Within the top 1% sequences ranked by PRISeq, the distribution of pairwise Hamming distances peaked at four nucleotides, with most pairwise Hamming distances falling between three and five positions, indicating that the set of the top 1% sequences was not dominated by closely related sequences. Most sequences differed from the wild-type sequences at two or three positions, and some differed at four positions. Clustering the top 1% sequences (1,292 sequences) yielded 186 sequence clusters, and even the largest cluster contained only 42 sequences, indicating broad dispersion across sequence space rather than convergence on a single dominant motif. Among these sequences, 137 sequences were experimentally active, corresponding to an EF_1%_ of 10.61%. Active sequences were distributed across multiple clusters, and five clusters consisted entirely of active sequences. Notably, three representative active sequences differing from the wild-type sequence at four positions were distributed across two clusters. Collectively, these results demonstrated that PRISeq identified diverse families of active sequences rather than merely enriching close variants of the wild-type sequence.

**Figure 3.**
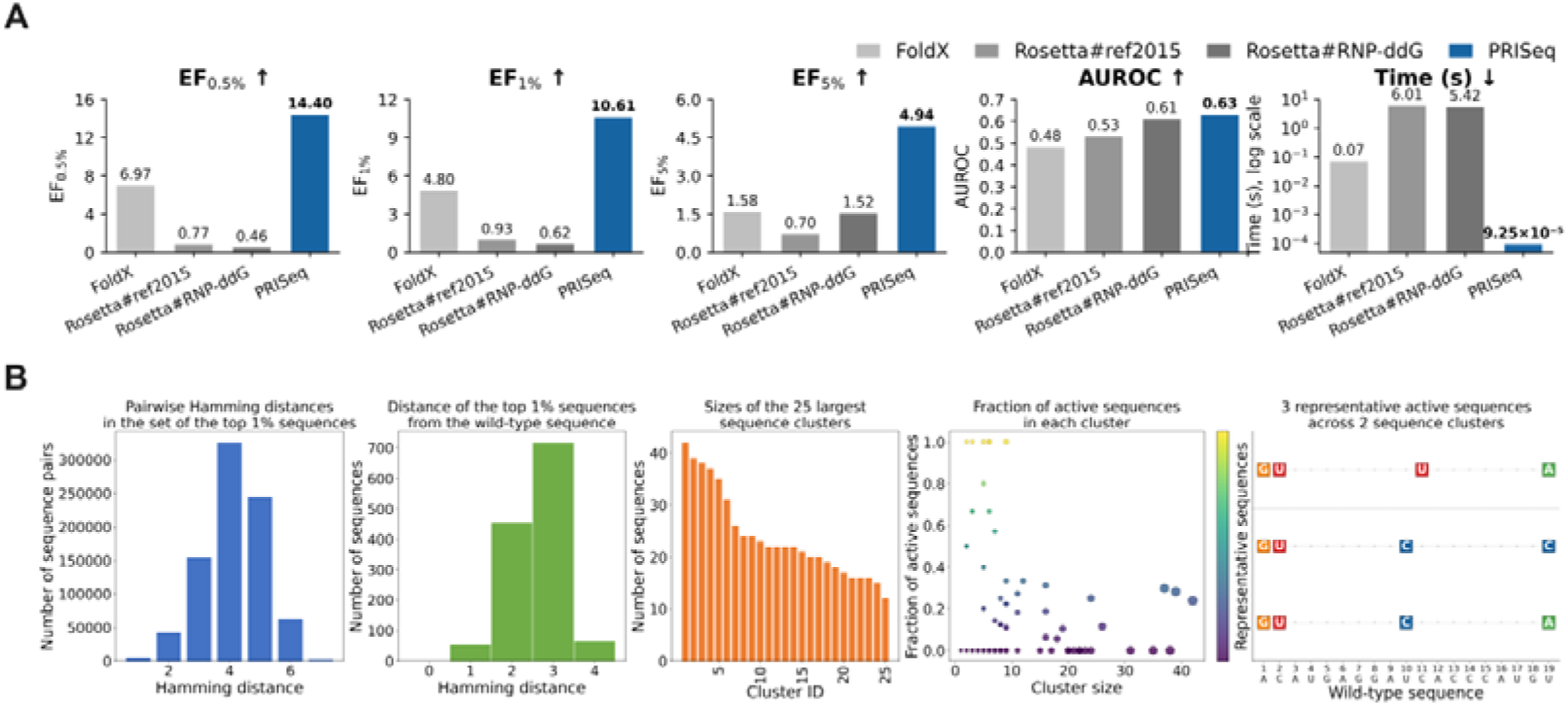
Performance of PRISeq in the virtual screening of a 129,248-member RNA library against the MS2 protein and the sequence diversity among the top 1% sequences ranked by PRISeq. (A) Comparison of PRISeq with FoldX, Rosetta#ref2015, and Rosetta#RNP-ddG using EF_0.5%_, EF_1%_, EF_5%_, AUROC, and the average computation time per sequence. Computation time is shown on a logarithmic scale. The reported times for the three SFs include energy evaluation but exclude mutation generation and energy minimization. The upward and downward arrows indicate that larger and smaller values correspond to the better performance, respectively. (B) From left to right, the panels show the distribution of pairwise Hamming distances, Hamming distances from the wild-type 19-nt RNA hairpin, the sizes of the 25 largest sequence clusters, the fraction of experimentally active sequences in each cluster, and the substitution patterns of three representative active sequences that differed from the wild-type sequence at four positions and were distributed across two clusters. In the scatter plot, point area represents the relative cluster size and color represents the fraction of active sequences. In the substitution map, letters and colored squares indicate substituted nucleotides (A, green; C, blue; G, orange; and U, red), whereas gray dots indicate positions identical to the wild-type sequence.

### PRIScore improves the success rates of protein-RNA structure predictions

In the PWM inference task, the experimentally determined crystal structures were used as input for PRISeq. However, in real-world drug discovery applications, crystal structures are often unavailable or incomplete. To address this limitation, several structure prediction methods were proposed, such as AlphaFold3, which demonstrated strong performance in protein-RNA complex structure prediction^24^. In this study, we assessed the performance of AlphaFold3 on the unbound protein-RNA docking benchmark, which comprised 188 crystal structures. For each structure, AlphaFold3 generated 100 predictions, which were ranked by their internal ranking scores. RMSD(R) and RMSD(P) indicated the RNA and protein RMSDs, respectively, between each prediction and the corresponding crystal structure. As shown in **Table 3**, AlphaFold3 achieved an average RMSD(R) of 4.74 Å and an average RMSD(P) of 1.40 Å for the top 1 predictions. A prediction with both RMSD(R) and RMSD(P) values below 6.0 Å was considered successful. The top N success rate was defined as the percentage of structures for which at least one of the top N predictions was successful. AlphaFold3 achieved a top 1 SR of 79.26% and a top 5 SR of 81.38% (**Table 3**). We then re-ranked the AlphaFold3 predictions using four SFs, including FoldX, Rosetta#ref2015, Rosetta#RNP-ddG, and PRIScore. Compared with AlphaFold3, PRIScore (top 1 SR = 81.91%, top 5 SR = 84.04%) improved the success rates, whereas FoldX (top 1 SR = 77.13%, top 5 SR = 80.32%), Rosetta#ref2015 (top 1 SR = 76.60%, top 5 SR = 79.79%), and Rosetta#RNP-ddG (top 1 SR = 77.13%, top 5 SR = 80.32%) did not. PRIScore also achieved the lowest RMSD(R) of 4.27 Å for the top 1 predictions, demonstrating its ability to prioritize the most accurate predictions.

**Table 3.** Benchmark evaluation of the methods for re-ranking protein-RNA structure predictions.

| Methods | Top 1 SR(%) ↑ | Top 5 SR(%) ↑ | RMSD(R)<sup>a</sup><br>↓ | RMSD(P) ↓ |
| --- | --- | --- | --- | --- |
| FoldX | 77.13 | 80.32 | 5.62 | **1.39** |
| Rosetta#ref2015 | 76.60 | 79.79 | 5.45 | 1.48 |
| Rosetta#RNP-ddG | 77.13 | 80.32 | 5.29 | 1.48 |
| AlphaFold3 | 79.26 | 81.38 | 4.74 | 1.40 |
| PRIScore | **81.91** | **84.04** | **4.27** | 1.41 |

<sup>a</sup>The units of RMSD(R) and RMSD(P) are both Å.

As shown in **Figure 4B**, the top 1 predictions were grouped into four RMSD bins: below 3 Å, between 3 Å and 6 Å, between 6 Å and 9 Å, and above 9 Å. For AlphaFold3 and all SFs, more than 90% of the top 1 predictions had an RMSD(P) below 3 Å, and more than 95% had an RMSD(P) below 6 Å. By contrast, approximately 65% of the top 1 predictions had an RMSD(R) below 3 Å and approximately 80% had an RMSD(R) below 6 Å. These results indicated that the success of protein-RNA complex structure prediction was determined primarily by the accuracy of the predicted RNA structure. Notably, PRIScore achieved not only the highest proportion (81.9%) of the top 1 predictions with an RMSD(R) below 6 Å but also the highest proportion (67.0%) with an RMSD(R) below 3 Å, indicating higher accuracy of the predicted RNA structure. Two representative cases (PDB IDs: 1C9S and 2C0B) are shown in **Figure 4C**. PDB 1C9S contains the transcription attenuation protein MtrB bound to an expanded GAGAU-repeat RNA with a regular alternation of right-handed (AGA) and left-handed (U5 and G1) twists^55^. The top 1 prediction ranked by AlphaFold3 showed opposite twist handedness, with the left-handed twists in PDB 1C9S predicted as right-handed and the right-handed twists predicted as left-handed. The top 1 predictions ranked by Rosetta#ref2015 and Rosetta#RNP-ddG both contained paired bases, whereas all bases were unpaired in PDB 1C9S. The top 1 prediction ranked by PRIScore was highly similar to PDB 1C9S, with an RMSD(R) of 1.33 Å, whereas the top 1 predictions ranked by the other methods all had RMSD(R) values above 24 Å. Regarding PDB 2C0B, which contains ribonuclease E bound to a 13-nt RNA^56^, the top 1 prediction ranked by PRIScore had an RMSD(R) of 0.94 Å, whereas the top 1 predictions ranked by the other methods all showed an outward extension of the RNA, with RMSD(R) values above 6 Å.

**Figure 4.**
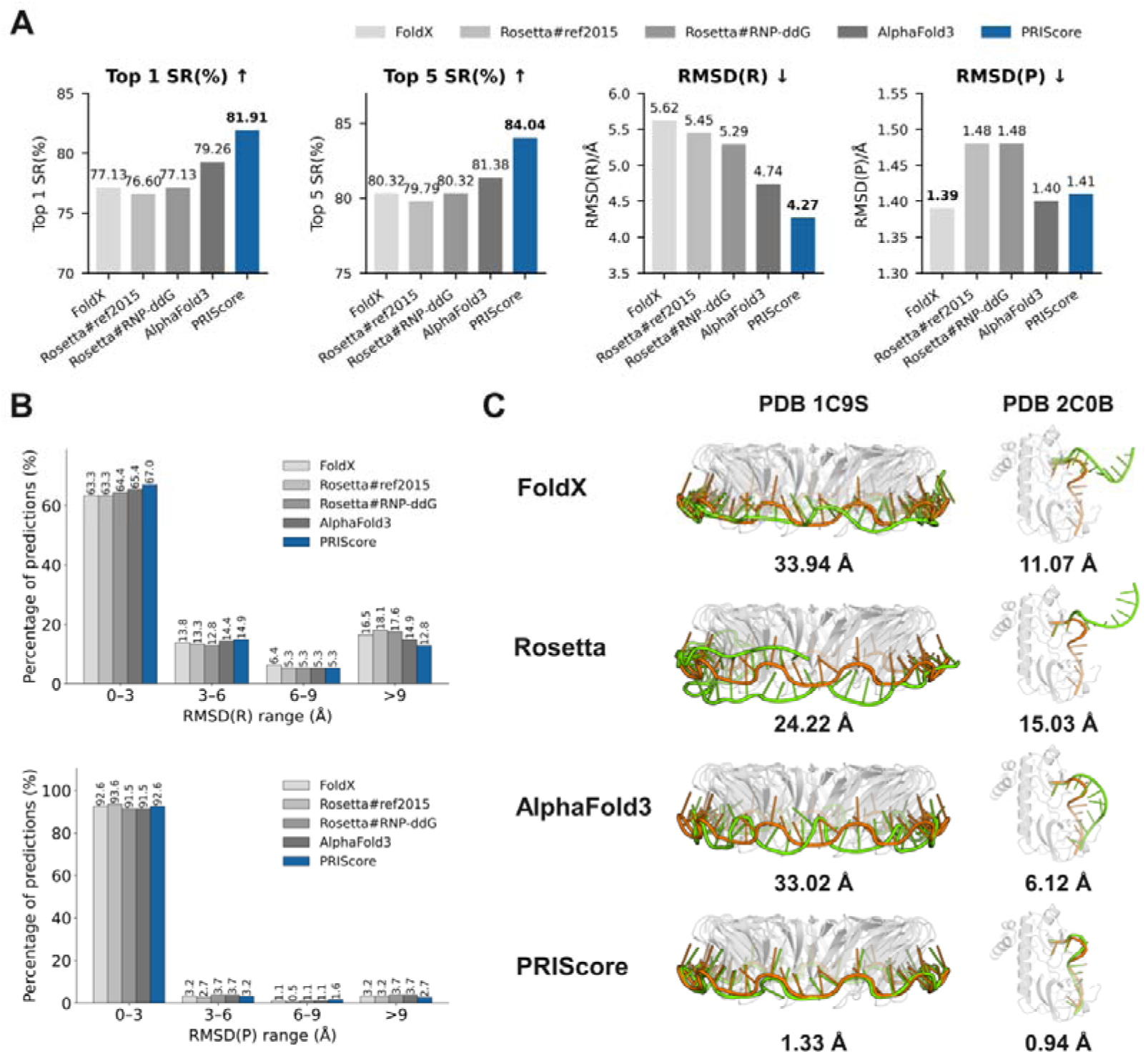
Benchmark evaluation of SFs for re-ranking protein-RNA structure predictions. (A) Comparison of PRIScore with FoldX, Rosetta#ref2015, Rosetta#RNP-ddG, and AlphaFold3. Performance is evaluated in terms of Top 1 SR, Top 5 SR, RMSD(R), and RMSD(P). The upward and downward arrows indicate that larger and smaller values correspond to the better performance, respectively. (B) Distributions of the top 1 predictions across four RMSD(R) (top) and RMSD(P) (bottom) bins. (C) Structural comparison of the top 1 predictions for two representative protein-RNA complexes, PDB 1C9S (left) and PDB 2C0B (right). The experimental and predicted RNA structures are shown in orange and green, respectively, and the protein structures are shown in grey. Rosetta denotes Rosetta#RNP-ddG. The corresponding RMSD(R) values are indicated below each structural comparison.

### PRIS screens the RNA aptamers against target proteins

We further evaluated PRIS on the more challenging task of screening RNA aptamers against two target proteins, NELF-E and GFP, as shown in **Figure 5A**. The binding affinities of 9,833 equal-length NELF-E aptamer (NELFapt) sequences and 1,875 equal-length GFP aptamer (GFPapt) sequences were experimentally measured^37^. However, no crystal structures were available for any of these aptamers, either in isolation or in complex with their respective target proteins. We used AlphaFold 3 to predict the structures of the wild-type protein-RNA aptamer complexes. For each complex, 100 predictions were generated and then re-ranked using the SFs. We compared the secondary structures of the top 1 predictions with the published secondary structures, which were supported by extensive mutational analysis^37^. Model performance was assessed using Precision, Recall, F1 score and BP distance. Precision measured the proportion of predicted base pairs that were also present in the published structure, whereas recall measured the proportion of published base pairs recovered in the prediction. The F1 score was calculated as the harmonic mean of precision and recall. BP distance was defined as the total number of base pairs present in one structure but absent from the other. Compared with AlphaFold3 (F1 score = 0.76, BP distance = 11), PRIScore (F1 score = 0.80, BP distance = 9) predicted more accurate RNA secondary structures, whereas other SFs did not. For example, the top 1 prediction ranked by PRIScore correctly reproduced the closed loop at the 5’-terminal region of NELFapt, whereas the top 1 prediction ranked by AlphaFold3 failed to recover this structural motif and instead displayed an open terminal region (**Figure 5B**). These results demonstrated that PRIScore identified more reliable predictions as inputs for subsequent aptamer screening using PRISeq.

**Figure 5.**
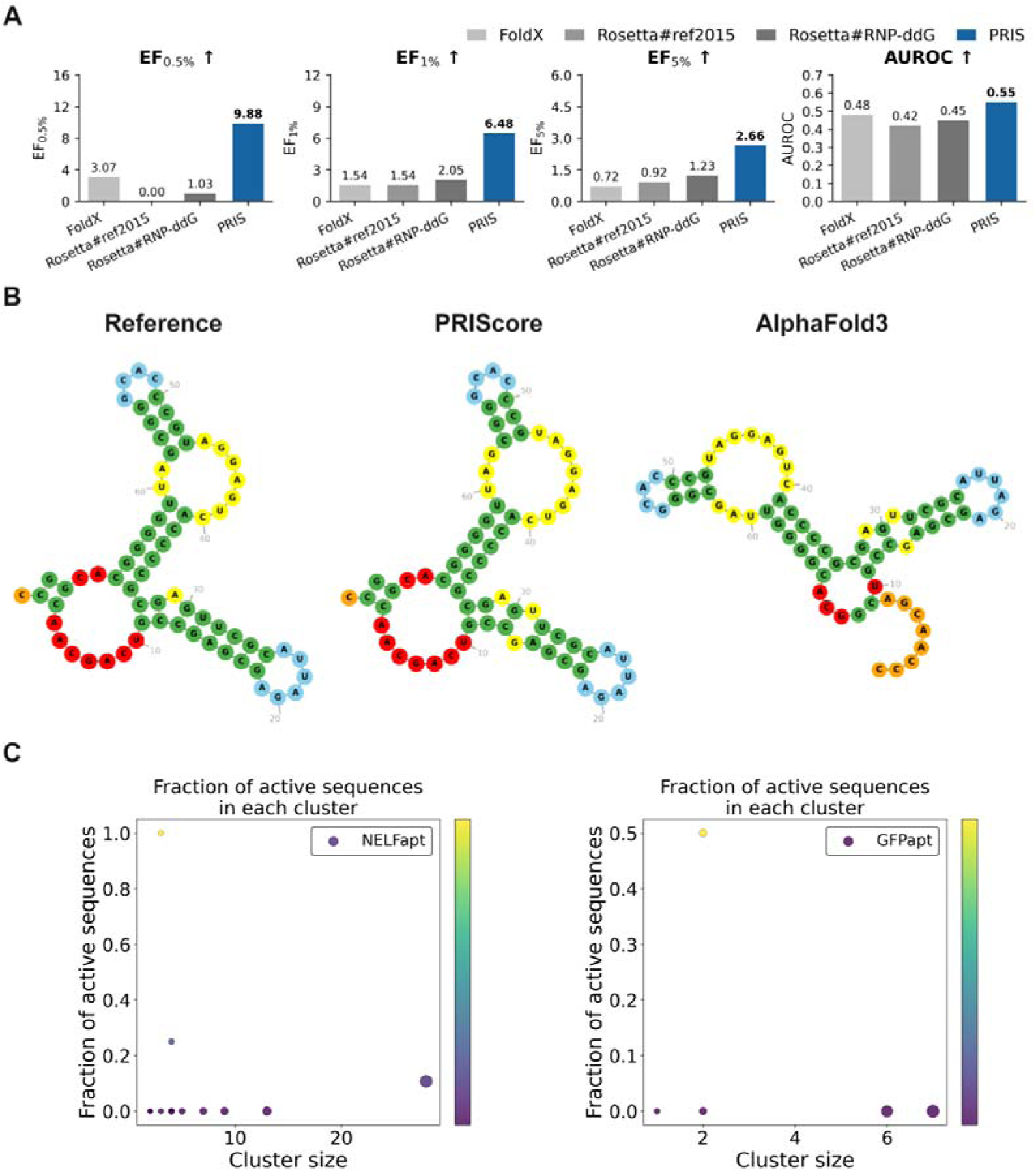
Evaluation of PRIS for RNA aptamer structure selection, virtual screening, and sequence diversity analysis. (A) Comparison of PRISeq with FoldX, Rosetta#ref2015, and Rosetta#RNP-ddG using EF_0.5%_, EF_1%_, EF_5%_, and AUROC. The upward and downward arrows indicate that larger and smaller values correspond to the better performance, respectively. (B) Reference secondary structure of NELFapt (left) and the secondary structures of the top 1 predictions ranked by PRIScore (middle) and AlphaFold3 (right). Stems, hairpin loops, internal loops, multiloops, and unpaired terminal regions are shown in green, blue, yellow, red, and orange, respectively. Base-pair connections are shown as red lines. The top 1 prediction ranked by PRIScore correctly reproduced the closed loop at the 5’-terminal region of NELFapt, whereas the top 1 prediction ranked by AlphaFold3 failed to recover this structural motif and instead displayed an open terminal region. (C) Fraction of experimentally active sequences in each cluster for NELFapt (left) and GFPapt (right). In the scatter plot, point area represents the relative cluster size and color represents the fraction of active sequences.

The top 1 predictions ranked by PRIScore were used as inputs for PRISeq. We compared PRISeq with three SFs, including FoldX, Rosetta#ref2015, and Rosetta#RNP-ddG. Regarding these SFs, the virtual screening workflow involved several steps, including mutation generation, energy minimization, and energy evaluation, as detailed in Materials and Methods, and the top 1 predictions ranked by AlphaFold3 were used as templates for mutation. Model performance was assessed using EF and AUROC. As shown in **Table 4**, PRIS consistently outperformed all baseline methods across the four evaluation metrics, achieving an EF_0.5%_ of 8.19, an EF_1%_ of 7.17, an EF_5%_ of 3.07, and an AUROC of 0.62 in the screening of NELFapt; an EF_0.5%_ of 11.57, an EF_1%_ of 5.79, an EF_5%_ of 2.24, and an AUROC of 0.48 in the screening of GFPapt; and an EF_0.5%_ of 9.88, an EF_1%_ of 6.48, an EF_5%_ of 2.66, and an AUROC of 0.55 on average. Compared with the best-performing baseline for each metric, PRISeq approximately tripled EF_0.5%_ and EF_1%_, approximately doubled EF_5%_, and increased the AUROC by 14.6% on average. These results highlighted the potential of PRIS for the efficient screening of RNA aptamers.

**Table 4.** Evaluation of the methods for RNA aptamer screening.

| Methods | EF<sub>0.5%</sub> ↑ | EF<sub>1%</sub> ↑ | EF<sub>5%</sub> ↑ | AUROC ↑ |
| --- | --- | --- | --- | --- |
| FoldX | 6.14/0.00/3.07 | 3.07/0.00/1.54 | 1.43/0.00/0.72 | 0.55/0.40/0.48 |
| Rosetta#ref2015 | 0.00/0.00/0.00 | 3.07/0.00/1.54 | 1.84/0.00/0.92 | 0.47/0.36/0.42 |
| Rosetta#RNP-ddG | 2.05/0.00/1.03 | 4.10/0.00/2.05 | 2.45/0.00/1.23 | 0.47/0.42/0.45 |
| PRIS | **8.19/11.57/9.88** | **7.17/5.79/6.48** | **3.07/2.24/2.66** | **0.62/0.48/0.55** |

PRIS also preserved sequence diversity in the screening of RNA aptamers. We examined the numbers of experimentally active sequences recovered among the top sequences ranked by PRISeq. For NELFapt, PRIS recovered 4, 7, and 15 active sequences among the top 49, 98, and 491 sequences, respectively, whereas random selection was expected to recover only 0.49, 0.98, and 4.91 sequences. Among the top 98 sequences, the seven active sequences were distributed across three sequence clusters: three belonged to the largest cluster, which contained 28 sequences; three formed a three-sequence cluster; and the remaining sequence belonged to a four-sequence cluster. Notably, four of the seven active sequences occurred in the two smaller clusters, indicating that PRISeq recovered active sequences from diverse sequence clusters rather than predominantly sampling a single cluster. For GFPapt, PRIS recovered 1, 1, and 2 active sequence among the top 9, 18, and 93 sequences, respectively, whereas random selection was expected to recover only 0.09, 0.18, and 0.93 sequences. Although the AUROC for GFPapt was 0.48, its high EF_0.5%_ of 11.57 indicated that PRIS still prioritized an experimentally activate sequence at the most stringent screening cutoff. Thus, the advantage of PRIS was pronounced when only a limited number of candidates could be experimentally evaluated.

## CONCLUSIONS

In this study, we present PRIS, a unified structure-based deep-learning framework. PRIS comprises two complementary components, PRISeq and PRIScore, which adopt the same feature extraction module but differ in their probability estimation modules. Their feature extractor combines an Anti-Symmetric Graph Attention Network with sparse k-Maximum Inner Product attention to capture long-range interactions in large graphs across multiple layers. Regarding probability estimation modules, PRISeq estimates probability distributions of nucleotide types, whereas PRIScore estimates probability distributions of residue-nucleotide distances. In the PRIS workflow, PRIScore is used to improve the selection of native-like protein-RNA predictions if the crystal structure is unavailable. The selected prediction is then input to PRISeq to infer RNA-binding preferences or screen candidate RNA sequences.

PRIS outperforms existing methods on independent PWM, docking and screening datasets. Compared with FoldX, Rosetta#ref2015, Rosetta#RNP-ddG, and NA-MPNN, PRIS achieves the lowest MAE of 0.75 on the PWM dataset. Compared with FoldX, Rosetta#ref2015, Rosetta#RNP-ddG, and AlphaFold3, PRIS achieves the highest top 1 SR of 81.91% on the docking dataset. Compared with FoldX, Rosetta#ref2015, and Rosetta#RNP-ddG, PRIS achieves the highest EF_0.5%_ of 14.40 in the screening of a 129,248-member RNA library within 11.95 s, and the highest EF_0.5%_ of 9.88 in the screening of RNA aptamers. PRIScore improves the secondary structure accuracy of predicted NELF-E aptamer, providing more reliable input for subsequent sequence screening using PRISeq.

By connecting structure selection with binding-preference inference, PRIS provides an efficient framework for interpreting protein-RNA recognition, accelerating large-scale RNA library screening, and aptamer design. It can also complement experimental selection by narrowing the sequence search space and proposing candidate sequences for subsequent validation, thereby reducing the experimental workload. The current evaluation covers a limited number of target proteins and ignores the target binding specificities of PBRs. Future work could evaluate PRIS on broader benchmarks, extend the framework to predict target binding specificity, thereby improving its robustness and extending its applicability to more diverse protein-RNA systems.

## Data and data availability

The datasets are available at https://doi.org/10.5281/zenodo.22651663. The code is available at https://github.com/roger-yh-zhao/PRIS.

## Author Contributions

T. J. H. and Y. K. and J. F. C. designed the research study. Y. H. Z. developed the method and wrote the code. Y. H. Z., J. H., J. K. W performed the analysis. Y. H. Z. and T. J. H. wrote the paper. All authors read and approved the manuscript.

## Acknowledgements

This work was financially supported by National Natural Science Foundation of China (22220102001, 92370130).

