## supplemental table 1,2,3 for "A Unified Structure-based Deep Learning Framework for High-Throughput Screening of Protein-Binding RNAs"

### Supporting information

**Table S1.** Node and edge features of nucleic acid graphs for PRISeq.

| Features | Size | Description |
|---|---|---|
| | | Nodes |
| self_distance | 11 | maximum distance within any atomic pair, distances between the atomic pairs O3'-C5', O3'-O5', O3'-P, C3'-O5', C3'-P, C4'-P, O3'-C1', O3'-O4', C5'-C1', C5'-C2' (multiplied by 0.1) |
| dihedral_angle | 16 | dihedral angles O5'(b)-P(b)-O3'-C3', P(b)-O3'-C3'-C4', O3'-C3'-C4'-C5', C3'-C4'-C5'-O5', C4'-C5'-O5'-P, C5'-O5'-P-O3'(f), P(f)-O3'-C3'-C2', O3'-C3'-C2'-C1', C3'-C2'-C1'-O4', C2'-C1'-O4'-C4', C1'-O4'-C4'-C5', O4'-C4'-C5'-O5', O3'-C3'-C4'-O4', C5'-C4'-C3'-C2', C3'-C4'-O4'-C1', C4'-C3'-C2'-C1' (multiplied by 0.01) |
| | | Edges |
| whether_connected | 1 | whether two nucleotides are connected |
| C5'_distance | 1 | distance between the atoms C5' of two nucleotides (multiplied by 0.1) |
| center_distance | 1 | distance between the centers of two nucleotides (multiplied by 0.1) |
| maximum_distance | 2 | maximum and minimum distances between two nucleotides (multiplied by 0.1) |

**Table S2.** Node and edge features of protein graphs for PRIS.

| Features | Size | Description |
|---|---|---|
| Nodes | | |
| type | 32 | residue type (["GLY", "ALA", "VAL", "LEU", "ILE", "PRO", "PHE", "TYR", "TRP", "SER", "THR", "CYS", "MET", "ASN", "GLN", "ASP", "GLU", "LYS", "ARG", "HIS", "MSE", "CSO", "PTR", "TPO", "KCX", "CSD", "SEP", "MLY", "PCA", "LLP", "metal", "other"]) with one hot encoding |
| self_distance | 5 | maximum and minimum distances within any atomic pair, distances between the atomic pairs CA-O, O-N, C-N (multiplied by 0.1) |
| dihedral_angle | 4 | dihedral angles phi, psi, omega, and chi1 (multiplied by 0.01) |
| ESM3_representation | 1536 | The hidden representation from ESM3 |
| Edges | | |
| whether_connected | 1 | whether two residues are connected |
| CA_distance | 1 | distance between the CA atoms of two residues (multiplied by 0.1) |
| center_distance | 1 | distance between the centers of two residues (multiplied by 0.1) |
| maximum_distance | 2 | maximum and minimum distances between two residues (multiplied by 0.1) |

**Table S3.** Evaluation of the methods for the prediction of RNA aptamer secondary structure.

| Methods | Precision ↑ | Recall ↑ | F1 score ↑ | BP distance ↓ |
|---|---|---|---|---|
| FoldX | 0.80/0.58/0.69 | 0.94/0.75/0.85 | 0.86/0.65/0.76 | 5/16/11 |
| Rosetta#ref2015 | 0.80/0.58/0.69 | 0.94/0.75/0.85 | 0.86/0.65/0.76 | 5/16/11 |
| Rosetta#RNP-ddG | 0.80/0.58/0.69 | 0.94/0.75/0.85 | 0.86/0.65/0.76 | 5/16/11 |
| AlphaFold3 | 0.80/0.58/0.69 | 0.94/0.75/0.85 | 0.86/0.65/0.76 | 5/16/11 |
| PRIScore | 0.90/0.58/0.74 | 1.00/0.75/0.88 | 0.95/0.65/0.80 | 2/16/9 |
